# Expression and engineering of unexplored PET degrading enzymes from *Microbispora, Nonomuraea, Micromonospora* genus

**DOI:** 10.1101/2023.04.17.537204

**Authors:** Elaine Tiong, Ying Sin Koo, Jiawu Bi, Lokanand Koduru, Winston Koh, Yee Hwee Lim, Fong Tian Wong

**Affiliations:** Institute of Molecular and Cell Biology (IMCB), Agency for Science, Technology and Research (A*STAR), 61 Biopolis Drive, Proteos #07-06, Singapore, 138673, Republic of Singapore; Institute of Sustainability for Chemicals, Energy and Environment (ISCE^2^), Agency for Science, Technology and Research (A*STAR), 8 Biomedical Grove, Neuros, #07-01, Singapore 138665, Republic of Singapore; Bioinformatics Institute (BII), Agency for Science, Technology and Research (A*STAR), 30 Biopolis Street, Singapore 138671, Republic of Singapore; Synthetic Biology Translational Research Program, Yong Loo Lin School of Medicine, National University of Singapore, 10 Medical Drive, Singapore 117597, Republic of Singapore

**Keywords:** PETase, PET degradation, *Microbispora*, *Nonomuraea*, *Micromonospora*, enzymes

## Abstract

Low recycling rates have resulted in the alarming rate of accumulation of a widely used plastic material, polyethylene terephthalate (PET). With the build-up of plastics in our environment, there is an urgent need to source for more sustainable solutions to process them. Biological methods such as enzyme-catalyzed PET recycling or bioprocessing are seen as a potential solution to this problem. Actinobacteria, known for producing enzymes involved in the degradation of complex organic molecules, are of particular interest due to their potential to produce PET degrading enzymes. The highly thermostable enzyme, leaf-branch compost cutinase (LCC) found in Actinobacteria is one such example. This work expands on the discovery and characterization of new PET degrading enzymes from *Microbispora, Nonomuraea*, and *Micromonospora* genus. Within this genus, we analyzed enzymes from the polyesterase-lipase-cutinase family, which have ∼60% similarity to LCC, where one of the enzymes was found to be capable of breaking down PET and BHET at 45-50 °C. Moreover, we were able to enhance the enzyme’s depolymerization rate through further engineering, resulting in a two-fold increase in activity.

**IMPORTANCE:** The proliferation of PET plastic waste poses a significant threat to human and environmental health, making it an issue of increasing concern. In response to this challenge, scientists are investigating eco-friendly approaches, such as bioprocessing and microbial factories, to sustainably manage the growing quantity of plastic waste in our ecosystem. Despite the existence of enzymes capable of degrading PET, their scarcity in nature limits their applicability. The objective of this study is to enhance our understanding of this group of enzymes by identifying and characterizing novel ones that can facilitate the breakdown of PET waste. This data will expand the enzymatic repertoire and provide valuable insights into the prerequisites for successful PET degradation.

## INTRODUCTION

Polyethylene terephthalate (PET) is amongst the most widely manufactured and utilized plastic material for consumer and industrial applications due to its favorable physiochemical properties and durability (1). With an estimated production of at least 1 million PET bottles every minute (2) and a global recycling rate of 9% (3), the rapid and large accumulation of non-biodegradable post-consumer PET waste either end up in landfills, terrestrial or aquatic environment. These PET that end up in the terrestrial or aquatic environment not only have a slow rate of decomposition, but can also subsequently enter the marine ecosystem as microplastics - posing risks to the environment and the health of both animals and humans (4, 5). Consequently, addressing the PET waste crisis is imperative for both environmental and health sustainability. Traditional solid waste treatment methods, such as landfill and incineration, have limitations in terms of secondary pollution and limited land resources. The main thermomechanical and chemical recycling methods are energy-intensive, requiring high temperatures, and can result in alterations to the properties of the PET plastic (6). Thus, there is a growing interest in less energy-intensive or “natural” methods for processing this material.

The discovery of the first PET degrading enzyme PETase from *Ideonella sakaiensis* (IsPETase) (7, 8) in 2016, gave hopes to a biological solution through enzymes and microbial factories for processing this inert plastic. Enzyme-catalyzed PET recycling or bioprocessing can proceed under mild reaction conditions, with minimal energy and chemical usage. This method of recycling is thus a more environmentally responsible alternative to petroleum-derived production processes. To improve PETase towards practical application, there has been numerous studies reporting more thermostable and active versions of the enzyme (9). Since then, various PET degradation enzymes have also been identified, including CALB lipase from *Candida antarctica* (10), cutinases from *Fusarium solani, Humicola insolens*, and *Thermobifida fusca* (11). According to the Plastics-Active Enzymes Database (PAZy), there are ∼3000 homologs of PET-active enzymes by profile hidden Markov models (12).

Actinobacteria is a diverse group of bacteria known to produce a wide range of enzymes that are utilized in various industrial and medical applications, such as antibiotic production and degradation of environmental contaminants (13, 14). In particular, the actinobacteria are of interest due to their ability to produce cutinases and hydrolases, enzymes involved in the degradation of complex organic molecules and synthesis of essential nutrients (14). One of the most outstanding and highly thermostable PET degrading enzymes found in the actinobacteria is the leaf-branch compost cutinase (LCC), which was mined from metagenomes (15). A recent report by Erickson et.al on another PET degrading enzyme within the actinobacteria family, further underscores the potential of actinobacteria as a source for PET depolymerization. Given the capabilities of actinobacteria in PET depolymerization, this study aim to explore other actinobacteria for new PET degrading enzymes (16).

In this work, we expand the enzymes for PET breakdown from actinobacteria through the discovery, engineering, and characterization of new and unexplored PET degrading enzymes from *Microbispora, Nonomuraea, Micromonospora* genus.

## RESULTS

### Genomic mining and *in silico* analysis of distant cutinase homologs

With reference to the reported LCC, potential hydrolases were obtained from the genera *Microbispora, Nonomuraea*, and *Micromonospora* through mining actinobacterial genomic sequences in public databases such as Genbank and Natural Products Discovery Center (NPDC) (17), as well as the Singapore-based Natural Organism Library (NOL) (18). Within these hits, we selected four representative sequences from NOL. Sequence alignment analysis between the four cutinases indicate sequence similarity (∼60-61 % homology to LCC, Fig. 1B), including the presence of the three active site triad residues for PET breakdown (Asp, Ser and His, Fig. 1A). Cysteine residues for disulfide bridge formation were also observed (Fig. S1). Search within the ESTHER database (19) also shows high similarity hits of these enzymes as part of the polyesterase-lipase-cutinase family; similar to many previously reported PET hydrolases. (Fig. S2). Absence of an extended loop and extra disulfide bond near the active site also classify these enzymes to be Type I (LCC) rather than Type II (IsPETase) (20). Structural alignment based on *in silico* AlphaFold models also proposed that their general structure is closely aligned to that of the LCC, including positions of the active site residues (Table S1, Fig. S1). Although the core active sites and general structures are conserved, electrostatic prediction of the surface indicated sequence divergence are mainly surface residues. This observation is consistent to conclusions drawn from an extensive study on PET breakdown enzymes (16).

**Figure 1.**
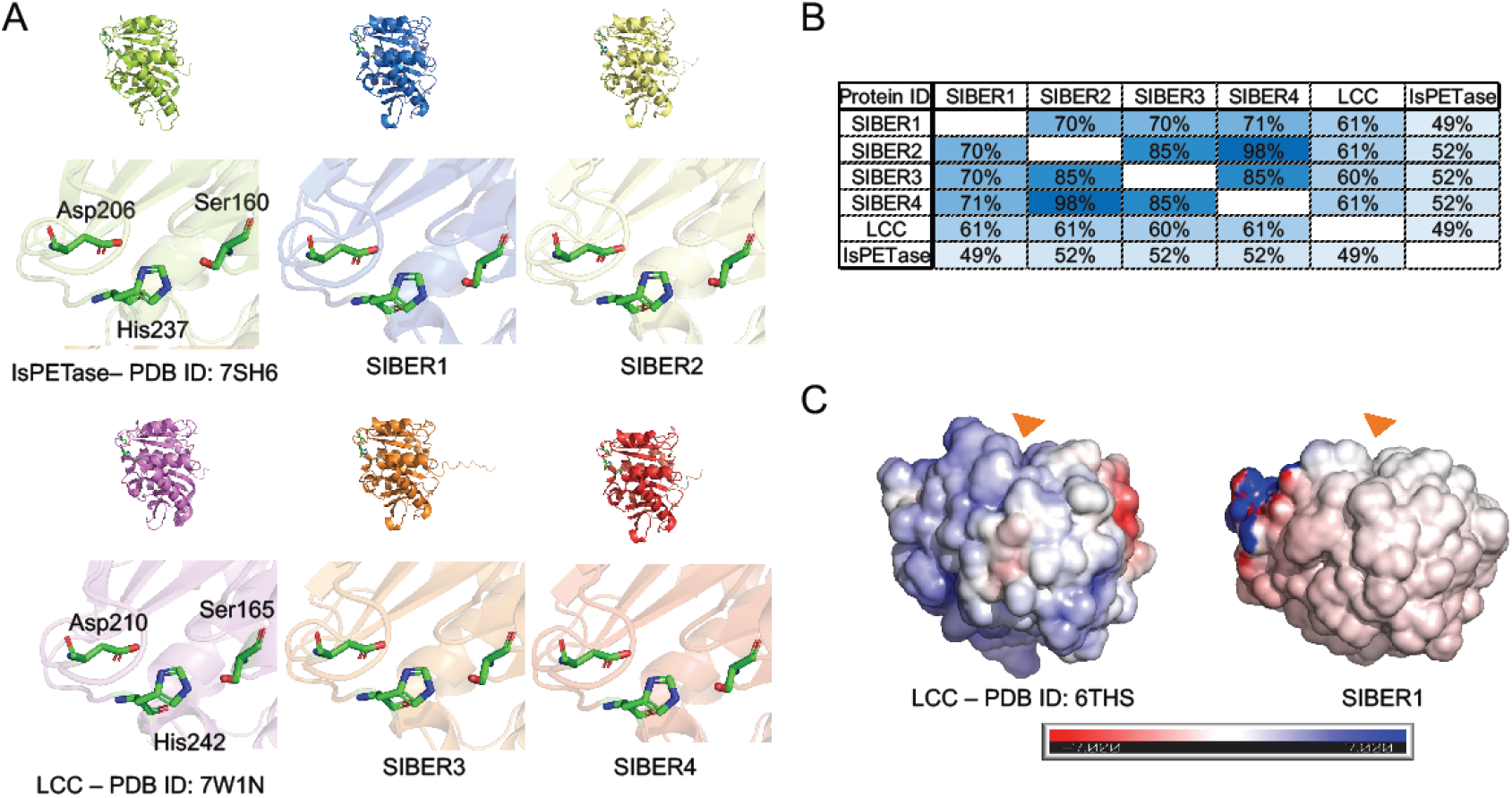
Genomic mining and *in silico* analysis of distant cutinase homologs. (A) Structural alignment of LCC and IsPETase against AlphaFold models of SIBER 1-4. Active sites residues are shown in inset below. RSMD of the *in-silico* models against 6THS range from 0.54 to 0.68 Å (Table S1). (B) Similarity scores between SIBER 1-4, LCC and IsPETase using BLASTP. (C) Electrostatic representative of solvent accessible surfaces of LCC and SIBER1 at pH7 (±7 kT/e, APBS-PDB2PQR (21))

### Optimizing heterologous expression

To investigate if the shortlisted sequences encode functional enzymes, there is a need to identify a suitable expression system. For heterologous expression of these enzymes in *Escherichia Coli* (*E. coli*), optimization was first performed for the protein expression constructs of SIBER 1-4. These were screened with various combinations of solubility tags for optimal expression (Fig. 2). Solubility aids which involved the Small Ubiquitin-like Modifier (SUMO (22)) and a 11 amino acid tag, NT11 (23), and Maltose Binding Protein (MBP) were investigated. Our optimization data showed varied expression levels across enzymes and tag combinations (Fig. 2). Overall, SIBER 1 constructs had significantly higher expression levels. The most productive constructs obtained from the distinct sequences were further scaled up and the resulting proteins were purified for characterization (SIBER 1-SIBER 171, SIBER 2-SIBER 196, SIBER 3-SIBER 207, SIBER 4-SIBER 228, Table S2-3, Fig. S3-S4). Protein yields ranged from 3-8 mg/L.

**Figure 2.**
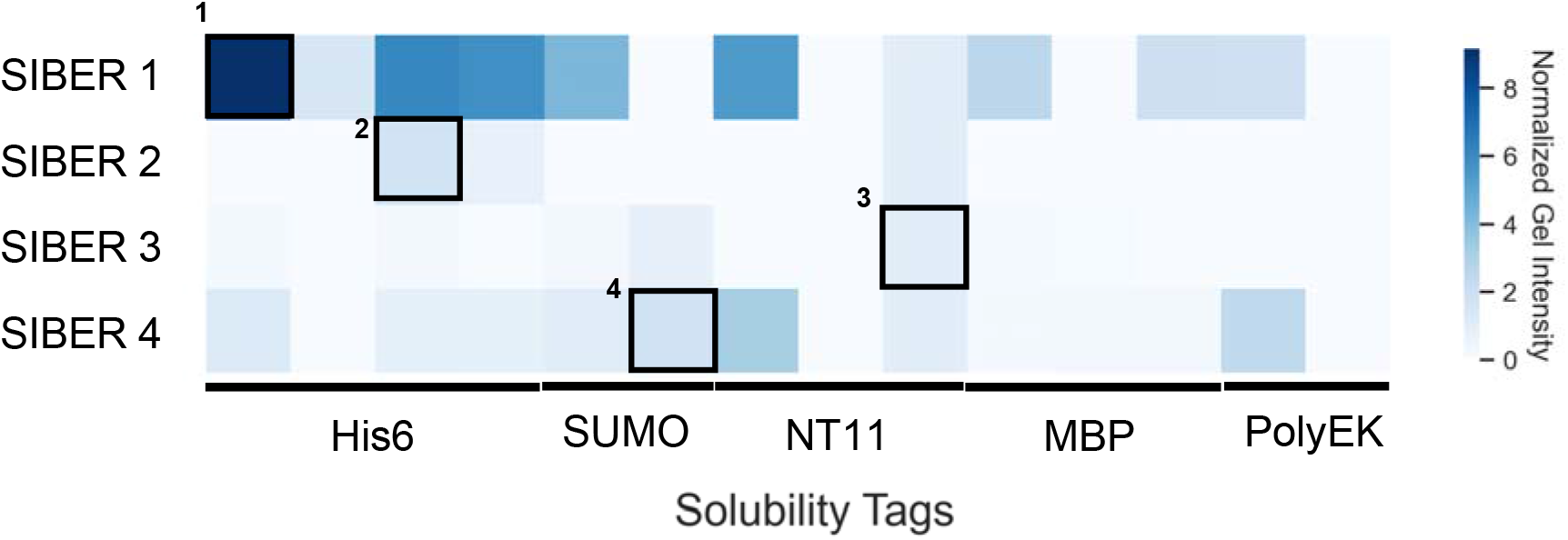
Protein expression optimization. This figure shows a heat map depicting expression levels of SIBER 1-4 genes fused with different solubility tags in *E. coli*. The final constructs were scaled up for characterization are annotated on the heat map; 1: SIBER 171, 2: SIBER 196, 3: SIBER 207, 4: SIBER 228. The proteins are His-tagged purified and ran on a protein gel. Quantification was made using ImageJ and proteins were normalized to proteins expressed with polyEK tags (last column).

### BHET and PET breakdown assays, characterization of SIBER 171, 196, 207, 228

To assess the functional activity of the enzymes, the purified enzymes (SIBER 171, SIBER 196, SIBER 207, SIBER 228) were subjected to various assays of different substrates (e.g., BHET, hcPET, lcPET). Under ambient temperature (25 °C) at pH 7, all the enzymes were able to hydrolyze BHET to various extents (Fig 3). The best mutant is SIBER 171 which resulted in ∼95% depolymerization by the end of 24 h. SIBER 171 was also functional up to 50 °C, whereas SIBER 196, SIBER 207 and SIBER 228 were only functional at 25 °C. Further characterization of SIBER 171 also showed that their activities are maintained across pH 6-8.

**Figure 3.**
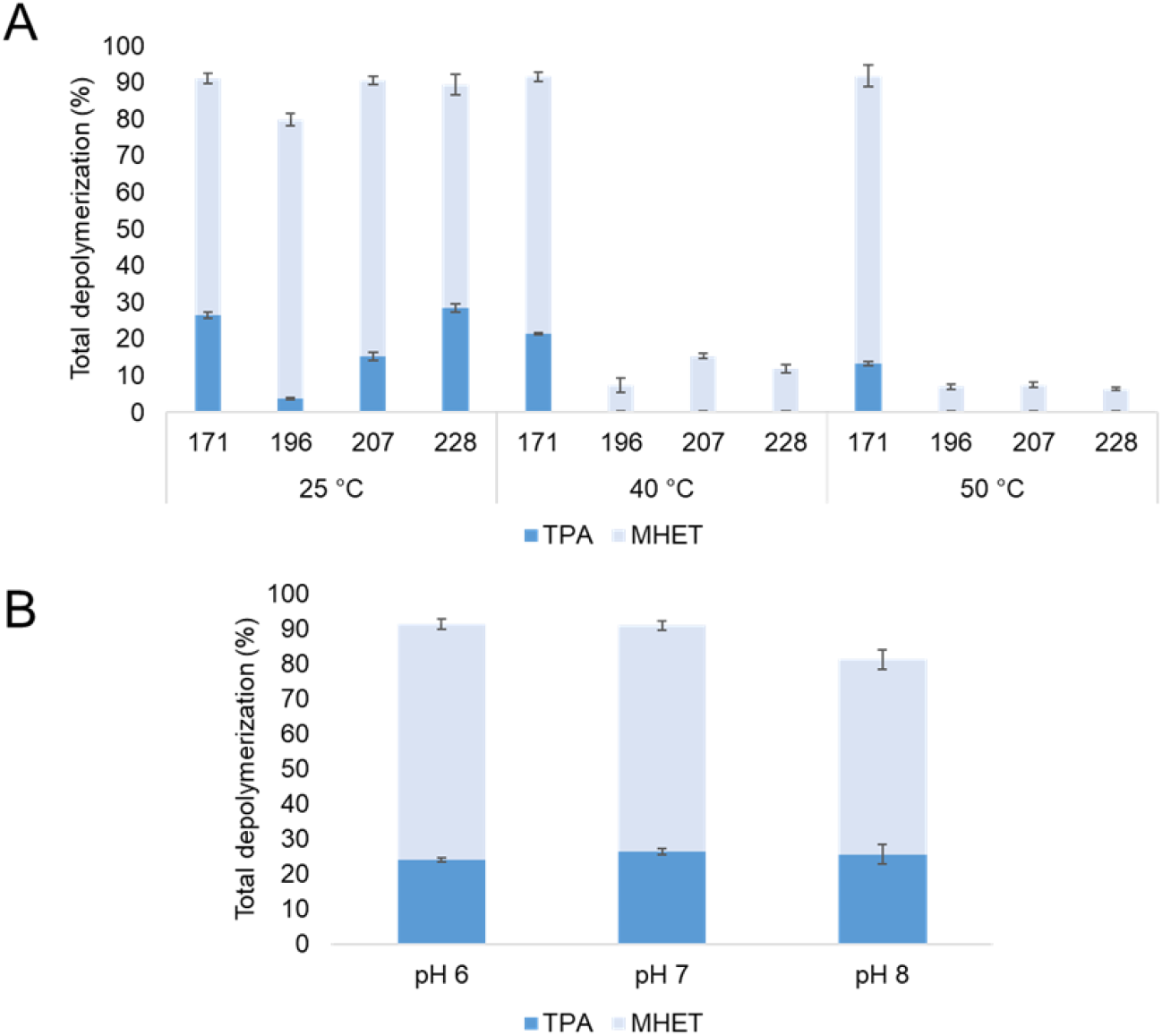
Characterization of SIBER 171, 196, 207 and 228. (A) Depolymerization of 5 mM BHET for 24 hours at various temperatures, pH 7. X axis are annotated as mutant numbering SIBER-. (B) Depolymerization of 5 mM BHET for 24 hours at various pH, 25°C, for SIBER 171.

Applying the reported optimized LCC conditions (70 °C, 48 h) for PET depolymerization to these enzymes did not show any PET breakdown, suggesting that the native enzymes are not tolerable to such high temperatures. Re-assaying these enzymes at lower temperatures (25 °C or 45 °C) over longer time (up to 3 weeks), we found that all five enzymes were able to assimilate high crystallinity PET (hcPET), though at a very slow rate (Fig. 4, S6). The best performing enzyme herein is again SIBER 171. Conversions mostly doubled in the second week when compared to the first week, indicating that the enzyme was still active. None of the variants are sufficiently thermostable to effectively break down high crystallinity (>35%) PET at 45 °C, though small amount of PET conversion was observed at 45 °C after 1 week for the enzymes SIBER 171, there was no further conversion after one week, suggesting that SIBER 171 might have been inactivated (Fig. 4, S6).

**Figure 4.**
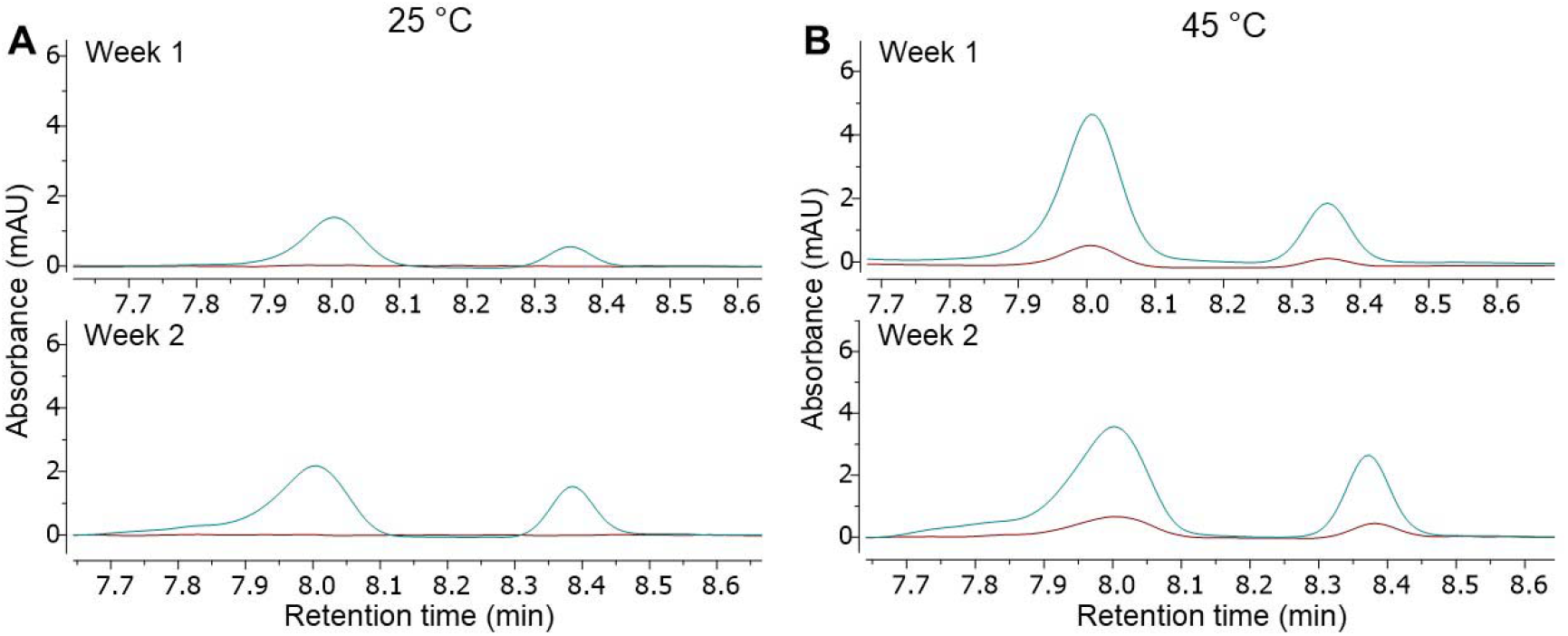
Depolymerization of high crystallinity PET (Goodfellow biaxially oriented film) with SIBER 171 (blue) and control (red) at (A) 25 °C over 2 weeks and (B) 45 °C over 2 weeks. The LC spectra extracted indicate peaks corresponding to TPA (RT 8 min and 8.3 –8.4 min, Fig. S8). Control reaction is the equivalent reaction setup in the absence of enzymes.

### Engineering of SIBER 171

To determine if similar optimizations made to LCC can be applied to improve SIBER 171, additional mutations were introduced, including the insertion of a disulfide bond for increased thermal stability (D238C/S283C, as numbered in (9)), F243I mutation to restore activity and N246M mutation for further optimization. Here, we made similar mutations for SIBER 171, resulting in SIBER 821 (SIBER 1-ICC) and SIBER 822 (SIBER 1-ICCM). As a result of these mutations, we were also able to obtain significantly higher yielding enzymes (30-40 mg/L, Fig. S5). Subsequently these purified enzymes were characterized for BHET breakdown at 50°C and PET breakdown at both 45 °C and 65 °C.

During BHET hydrolysis (50 °C, pH 7), we observed detrimental effects of ICC mutations which are however restored with N246M mutation (Fig. 5a). Under these conditions, there are minimal differences between native and engineered enzymes. However, using low crystallinity PET (lcPET) substrates at 45 °C, there appears to be significant improvement of SIBER 822 over the native enzymes over one week (Fig. 5). At 65 °C, no PET depolymerization was observed for both enzymes. Our observations suggest that, despite not being enough to increase its working temperature, the addition of a di-sulfide bridge to SIBER 1 resulted in potentially better stability at 45°C over 1 week when compared to its native enzyme. Although this is lower than optimized LCC (9) which yield PET depolymerization of 40% after 48 hours (Fig. S7), it is still encouraging to note that SIBER 822 is significantly more active than IsPETase double mutant (24) (∼0.08%-0.09% at 48 hours, Table S4). We anticipate that the further improvement of the enzyme towards practical applications will necessitate the implementation of additional engineering techniques, such as directed evolution and computational design.

**Figure 5.**
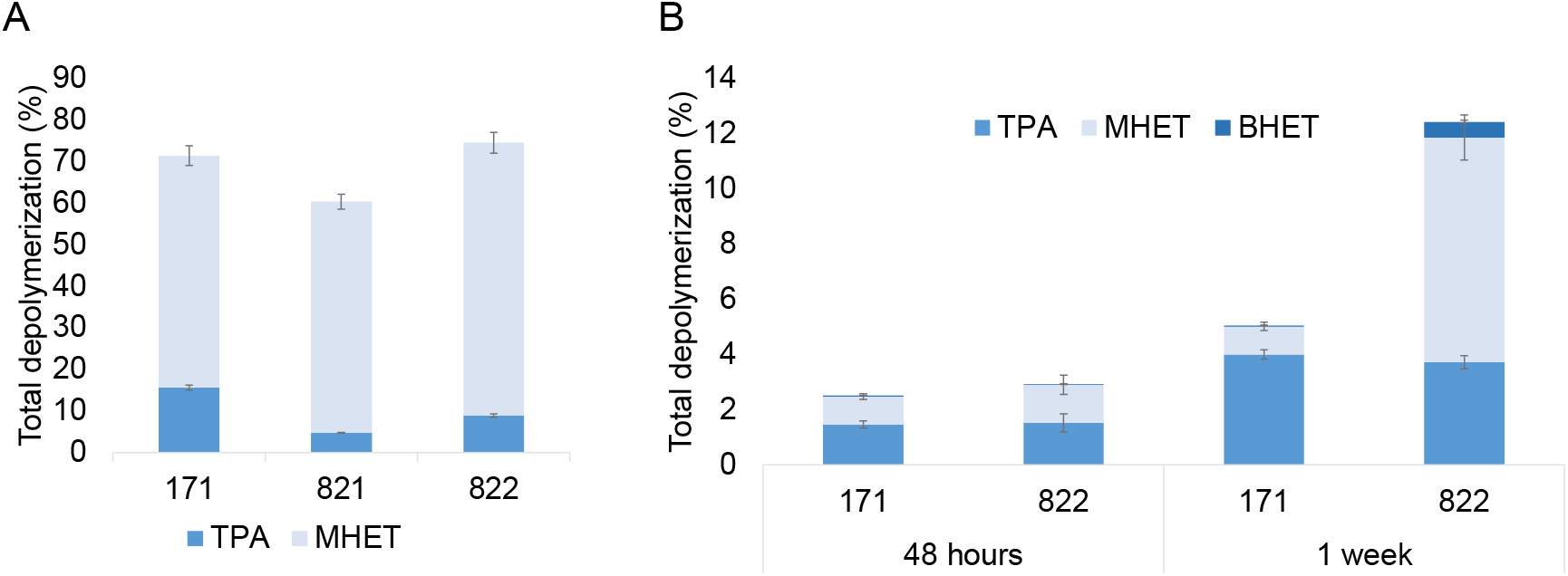
Engineering SIBER 171. (A) Depolymerization of 5 mM BHET for 50 °C, pH 7, 24 hours. X axis are annotated as mutant numbering SIBER-. (B) Depolymerization of low crystallinity PET (Goodfellow amorphous film) at 45 °C over 48 hours and 1 week. X axis (mutant numbering SIBER-). X axis are annotated as mutant numbering SIBER-.

## DISCUSSION

This study aims to deepen the understanding of PET-degrading enzymes by identifying and describing new ones that can aid in PET waste biodegradation. Although some PET-degrading enzymes are known, their rarity restricts the utilization of it. Even with the addition of 34 more recently characterized PET degrading enzymes from the actinobacteria family (16), this still adds up to only a few dozen of verified PET-active enzymes (12). Exploring a new genus space, we found hydrolases that have varying specificities towards substrates BHET, MHET, and even PET. Within representative four sequences, we have uncovered and characterized a new thermostable PETase within the genus of *Microbispora, Nonomuraea*, and *Micromonospora* genus. Furthermore, despite sharing only 60% homology with LCC, we discovered that incorporating similar mutations previously used in the engineering of LCC (9) could improve SIBER 171’s activity and stability. The engineered enzyme (SIBER 822) exhibited over two-fold improvement for lcPET depolymerization compared to its wild type.

To date, there are over 2000 known PET homologs and several enzyme engineering studies have proposed targeted mutations to improve thermostability, activity, and accessibility to PET. The results of this work raise questions about the factors that determine a PETase and the relationship between thermostability and PET degradation efficiency, especially in light of the impact of the substrate’s physical properties (25), such as crystallinity. Among the four enzymes, there were notable differences and correlations in solubility and expression yields and activity. Here, among the SIBER 1-4 genes, SIBER 1 constructs had consistent higher protein expression yields. After engineering of SIBER 171, we also observed significant expression yield increase in engineered SIBER 822. This is consistent with the discussions of correlations between protein folding and protein activity, where the thermodynamics of the sequences may play a role in proper and accurate folding (26). Beyond stability, further investigation on the mechanism of PET degradation is required to gain a deeper understanding on the potential to design and develop new and highly efficient PETases. The understanding on enzyme-plastic interactions would allow PETases to be applied to other plastics and feedstocks.

## MATERIALS AND METHODS

All solvents used in product analysis (acetonitrile, formic acid, dimethylformamide) were purchased from commercial suppliers. Bis (2-hydroxyethyl) terephthalate (BHET), terephthalic acid (TPA), potassium phosphate (NaH_2_PO_4_ and Na_2_HPO_4_), and 4-bromobenzoic acid were obtained from Sigma-Aldrich. High crystallinity PET film (ES301250) and low crystallinity PET film (ES301445) were obtained from Goodfellow and was ground into powder.

### Construction of plasmids optimized for *E. coli* expression

Codon-optimized DNA sequences obtained from *E. coli* were synthesized from Twist Biosciences. These inserts were assembled via golden gate into pET28a (+), cloned with *E. coli* Omnimax competent cells. *E. coli* XJb (DE3) autolysis competent cells (Zymo Research) was subsequently used for the transformation of these 6xHis tag constructs for protein expression. Table S2 shows the amino acid sequences of the gene used and the solubility tags used are shown in Table S3.

### High throughput screening of heterologous expression constructs

1 mL overnight autoinduction media (Merck) was used for the inoculation of single colonies obtained from the transformation, in which 50 μg/mL kanamycin, 1.5 M L-arabinose and 0.5 M magnesium chloride was also added. A single freeze thaw cycle in lysis buffer (50 mM sodium phosphate buffer, 300 mM sodium chloride, 10 mM Imidazole and 0.03 % TritonX-100), from the cell pellet harvested allowed the cell to release its embedded protein. To capture these His-tagged proteins, Ni resin (PureCube, Cube Biotech) was used. These proteins were subsequently eluted in 50 mM sodium phosphate buffer pH 7.0, 300 mM sodium chloride and 500 mM imidazole. Quantification of the protein elutes on a protein gel was performed by ImageJ to compare protein yields across the different constructs. Intensity of each band obtained was normalized to the intensity of the reference band on the standard ladder used, as well as molecular weight of each protein. This densitometric comparison was used to determine the best yielding constructs which were then scaled up for purification and subsequent characterization.

### Protein expression and purification

A starter culture of 5 mL grown in LB Broth containing 50 μg/mL kanamycin was prepared, where a single colony from the transformation was inoculated overnight at 37 °C. The starter culture was diluted 200-fold in a fresh 1 L LB Broth containing 50 μg/mL kanamycin, 1.5 M L-arabinose and 0.5 M Magnesium chloride. Subcultures were grown at 37 °C until the optical density at 600 nm (OD600) reached ∼0.4 – 0.5. 100 μM isopropyl β-D-1-thiogalactopyranoside (IPTG) was then added to induce the expression of proteins. Following which, the cultures were incubated overnight at 16 °C for 18 to 20 hours. Cells were harvested from this culture using a centrifuge (15 minutes, 8000 *g*) maintained at 4 °C, before resuspending and freezing the resulting pellet at -80 °C in 10 mL of lysis buffer containing 50 mM sodium phosphate buffer pH 7.0, 300 mM sodium chloride, 10 mM Imidazole and 0.03 % TritonX-100. To ensure protein stability, subsequent purification steps were also carried out at 4 °C.

The frozen pellet was thawed with the addition of 10mL lysis buffer and sonicated to release proteins from the cells. The supernatant obtained from centrifugation of lysates at 13,500xg was loaded on Ni resin and incubated for 1 hour to capture the His-tagged proteins. 20mL of 50mM sodium phosphate buffer, 300mM sodium chloride and 50mM imidazole was then used to wash the resin, before eluting the bound proteins with 50mM sodium phosphate buffer pH 7.0, 300mM sodium chloride and 500mM imidazole. Lastly, the proteins were exchanged into 50 mM sodium phosphate pH 7.0 with 10% glycerol to allow for long-term storage at -80 °C. Subsequent BHET and PET degradation studies were carried out with these purified proteins.

### BHET depolymerization assay

Dimethylformamide was used to dissolve 1 M BHET stock solution. BHET stock solution (2.5 μL, 5 mM) was pipetted into a 2 mL glass vial containing 500 μL of 100 mM potassium phosphate buffer (pH 7) and 1.67 μM purified protein. The glass vial was tightly capped, and the reaction mixture was incubated at 25 °C for 24 hours in an Eppendorf ThermoMixer^®^. After 24 hours, the reaction is quenched with 500 μL of methanol. The mixture was transferred into a 10 mL centrifuge tube, the vial was washed with 4 mL of buffer/methanol (1:1, v/v) and the contents were transferred to the centrifuge tube. This is followed by the addition of 5 mL 0.5 mM 4-bromobenzoic acid in buffer/methanol (1:1, v/v) as an internal standard. The reaction mixture was sonicated, and an aliquot was filtered with a 0.2 μm syringe filter and analyzed via UHPLC-MS.

### Calibration of BHET and TPA

BHET and TPA stock solutions (1 mM) were prepared by dissolving the solids in 5 mL of in buffer/methanol (1:1, v/v), followed by the addition of 5 mL 0.5 mM 4-bromobenzoic acid in buffer/methanol (1:1, v/v) as an internal standard. 6 concentrations ranging from 0.05 mM to 1 mM were prepared from the stock solution and analyzed via UPHLC-MS. A calibration was plotted with the molar ratio against area ratio. LC spectrum of TPA (Fig. S8) revealed 2 peaks (RT 8 min and RT 8.4 min) and the LC-MS spectrum extracted from both peaks correspond to TPA [M-H]^-^ ion m/z 165 (Fig. S8). The area of both peaks was taking into consideration in the plotting of calibration curve of TPA (Fig. S8).

### Calibration of MHET

MHET (1 M) was prepared by dissolving the solids in dimethylformamide. 6 concentrations ranging from 0.03 mM to 0.18 mM were prepared from the MHET stock solution, topped up with 0.5 mM 4-bromobenzoic acid internal standard and analyzed via UPHLC-MS. A calibration was plotted with the molar ratio against area ratio.

### PET depolymerization assay

PET powder (2 mg, 20 mM) was weighed into a 2 mL glass vial and fully submerged in 500 μL of 100 mM potassium phosphate buffer (pH 7) with 1.67 μM purified protein. The glass vial was tightly capped, and the reaction mixture was incubated at 500 rpm and 25-50°C for 48 hours to up to 28 days in an Eppendorf ThermoMixer^®^. The reaction is quenched with 500 μL of methanol. The mixture was transferred into a 10 mL centrifuge tube, the vial was washed with 4 mL of buffer/methanol (1:1, v/v) and the contents were transferred to the centrifuge tube. This is followed by the addition of 5 mL 0.5 mM 4-bromobenzoic acid in buffer/methanol (1:1, v/v) as an internal standard. The reaction mixture was sonicated, and an aliquot was filtered with a 0.2 μm syringe filter and analyzed via UHPLC-MS.

## Abbreviations

lcPET: Low crystallinity poly(ethylene terephthalate)
hcPET: High crystallinity poly(ethylene terephthalate)
PET: Poly(ethylene terephthalate)
TPA: Terephthalic acid
BHET: Bis(2-hydroxyethyl) terephthalic acid
MHET: Mono(2-hydroxyethyl) terephthalic acid
LCC: Leaf-branch compost cutinase
IsPETase: ldeonella sakaiensis PETase
SUMO: Small Ubiquitin-like Modifier
MBP: Maltose Binding Protein

## ACKNOWLEDGMENTS

This research is supported by Agency for Science, Technology and Research, Singapore, A*STAR <C211917006> and <C211917003>. J.B. acknowledges A*STAR Graduate Academy (A*GA) for his scholarship funding. Sequences derived for this work were mined from the in-house A*STAR National Organism Library (NOL) and public Genbank database. We gratefully acknowledge Dr. Siew Bee Ng (SIFBI), her team and Elena Heng for their contribution to the sequencing of the strains from NOL. A patent application has been filed for this work.

